# Cocaine regulation of *Nr4a1* chromatin bivalency and mRNA in male and female mice

**DOI:** 10.1101/2022.04.22.489203

**Authors:** Delaney K. Fischer, Keegan S. Krick, Chloe Han, Morgan Woolf, Elizabeth A. Heller

## Abstract

**BACKGROUND:** Cocaine epigenetically regulates gene expression via changes in histone post-translational modifications (HPTMs). We previously found that the immediate early gene *Nr4a1* is epigenetically activated by cocaine in mouse brain reward regions. HPTMs act combinatorically, yet few studies examine multiple HPTMs at a single gene. Bivalent gene promoters are simultaneously enriched in both activating (H3K4me3 (K4)) and repressive (H3K27me3 (K27)) HPTMs. As such, bivalent genes are lowly expressed but poised for activity-dependent gene regulation. In the current study, we defined regulation of K4&K27 bivalency at *Nr4a1* following cocaine treatment in male and female mice. The inclusion of female mice can shed light on the epidemiological relevance of sex to cocaine use disorder.

**METHODS:** Male and female mice were injected with saline or cocaine (i.p. 20mg/kg). We applied sequential chromatin immunoprecipitation and qPCR to define *Nr4a1* bivalency and expression in striatum (STR), prefrontal cortex (PFC), and hippocampus (HPC). Pearson’s correlation matrices quantified relationships within each brain region across treatment conditions for each sex.

**RESULTS:** We defined K4&K27 bivalency at the *Nr4a1* promoter in all three brain regions, in both sexes. In female STR, cocaine increased *Nr4a1* mRNA, coupled to maintenance of *Nr4a1* K4&K27 bivalency. In male STR, cocaine enriched repressive H3K27me3 and K4&K27 bivalency at *Nr4a1* and failed to increase *Nr4a1* mRNA. Furthermore, cocaine epigenetically regulated a putative NR4A1 target, *Cartpt*, in male PFC.

**CONCLUSION:** This study defined the epigenetic regulation of *Nr4a1* in reward brain regions in male and female mice. Cocaine treatment in female mice increased *Nr4a1* mRNA in STR, but there was no change in *Nr4a1* H3K27me3 or K4&K27 promoter bivalency. Following cocaine treatment in male mice, *Nr4a1* mRNA did not change in STR, HPC, or PFC, and *Nr4a1* H3K27me3 and K4&K27 promoter bivalency increased in the STR.

## INTRODUCTION

Despite decades of research, cocaine use disorder remains a global health problem for which there is currently no FDA approved treatment. On a molecular level, cocaine regulates gene expression in part through changes in histone post-translational modifications (HPTMs). Such changes can be transient or persist across long periods of abstinence (1–8). HPTMs tune gene expression by altering chromatin accessibility and recruiting transcriptional machinery (3). Acute and chronic cocaine impact HPTMs in multiple brain reward regions (7,9,10), such as the striatum (STR) (8,11), hippocampus (HPC) (12–14), and prefrontal cortex (PFC) (15,16). However, the complexities of chromatin remodeling following cocaine has yet to be explored in both sexes.

The histone code has historically proposed that engagement of individual HPTMs at a chosen locus can induce changes in protein recruitment and impact gene expression. However, the code also posits that HPTMs can be interdependent and/or work in tandem to impact gene expression (17,18). Thus, the histone code suggests that chromatin accessibility and, correspondingly, gene expression are largely determined not only by the concentration of single HPTMs at a given gene promoter, but by the combination of HPTMs at a given locus (17).

While many studies demonstrate genome-wide regulation of individual HPTMs following cocaine exposure (3,19–21), the literature lacks investigation of how cocaine impacts co-localization of HPTMs. Bivalent gene promoters are comprised of co-localized activating (H3K4me3 (K4)) and repressive (H3K27me3 (K27)) HPTMs (22) at a single promoter. Equivalent enrichment of H3K4me3 and H3K27me3 results in a poised state of gene expression, and can be induced by the addition and co-localization of either HPTM. Alternatively, bivalency can be ‘resolved’ to an active or repressive state by removal of H3K27me3 or H3K4me3, respectively (22). While limited studies have investigated bivalency in the brain (23–25), none to date have examined bivalency following cocaine exposure or sex, specifically.

We, and others, recently discovered that cocaine exposure activates expression of nuclear receptor subfamily 4 group A member 1 (*Nr4a1*) in the nucleus accumbens (NAc), a part of the STR (2,26,27). Studies show that activation of *Nr4a1* in the NAc attenuates cocaine self-administration (26), cocaine conditioned place preference (26), cocaine sensitization (28), and amphetamine-induced locomotion (29). *Nr4a1* functions as a transcription factor at target genes involved in drug reward, such as *Cocaine- and amphetamine-regulated transcript* (*Cartpt), Tyrosine Hydroxylase (Th), solute carrier family 6 member 3* (DAT production), and Brain-derived neurotrophic factor (*Bdnf)* (26,30,31), as well as plasticity in reward-related brain regions (28,32). However, the precise function of HPTMs in cocaine-induced *Nr4a1* expression has yet to be elucidated. Additionally, there is no data on the regulation of *Nr4a1* where sex is a defined variable. Accordingly, we investigated K4&K27 bivalency in the cocaine regulation of *Nr4a1* expression in both male and female mice.

In this study, we utilized single sample sequencing (S3EQ) (33) and sequential ChIP (23,34) to investigate the sex-specific transcriptomic and epigenomic profiles of *Nr4a1* from three different brain regions. We also utilized Pearson’s correlation matrices to enable us to capture the individual variability in biological changes induced by cocaine exposure in males and females.

## Methods and Materials

### Animals

Male and female mice on the C57BL/6J background were used in this study. Mice were housed under a 12-hour light-dark cycle at 23 °C with access to food and water *ad libitum*. Note that in compliance with ethical standards to minimize the use of mice, the mice used in this study were cre-negative offspring of *R26-CAG-LSL-Sun1-sfGFP;*A2a-cre and *LSL-Sun1-sfGFP*;Drd1-cre, generated for a separate study. All animal procedures were conducted in accordance with the National Institutes of Health Guidelines as well as the Association for Assessment and Accreditation of Laboratory Animal Care. Ethical and experimental considerations were approved by the Institutional Animal Care and Use Committee of The University of Pennsylvania.

### Investigator-administered cocaine

Male and female mice were handled daily for three days prior to cocaine administration. Following handling, mice were given a daily cocaine hydrochloride intraperitoneal injection (20 mg/kg dissolved in 0.9% saline) or 0.9% saline injection for 10 days (26). Male and female mice were injected with cocaine at the same time. Mice were sacrificed one day after the final injection. STR, HPC, and PFC tissue was immediately collected from each animal **(Figure 1A, B)** and stored at −80°C until processing.

**Figure 1.**
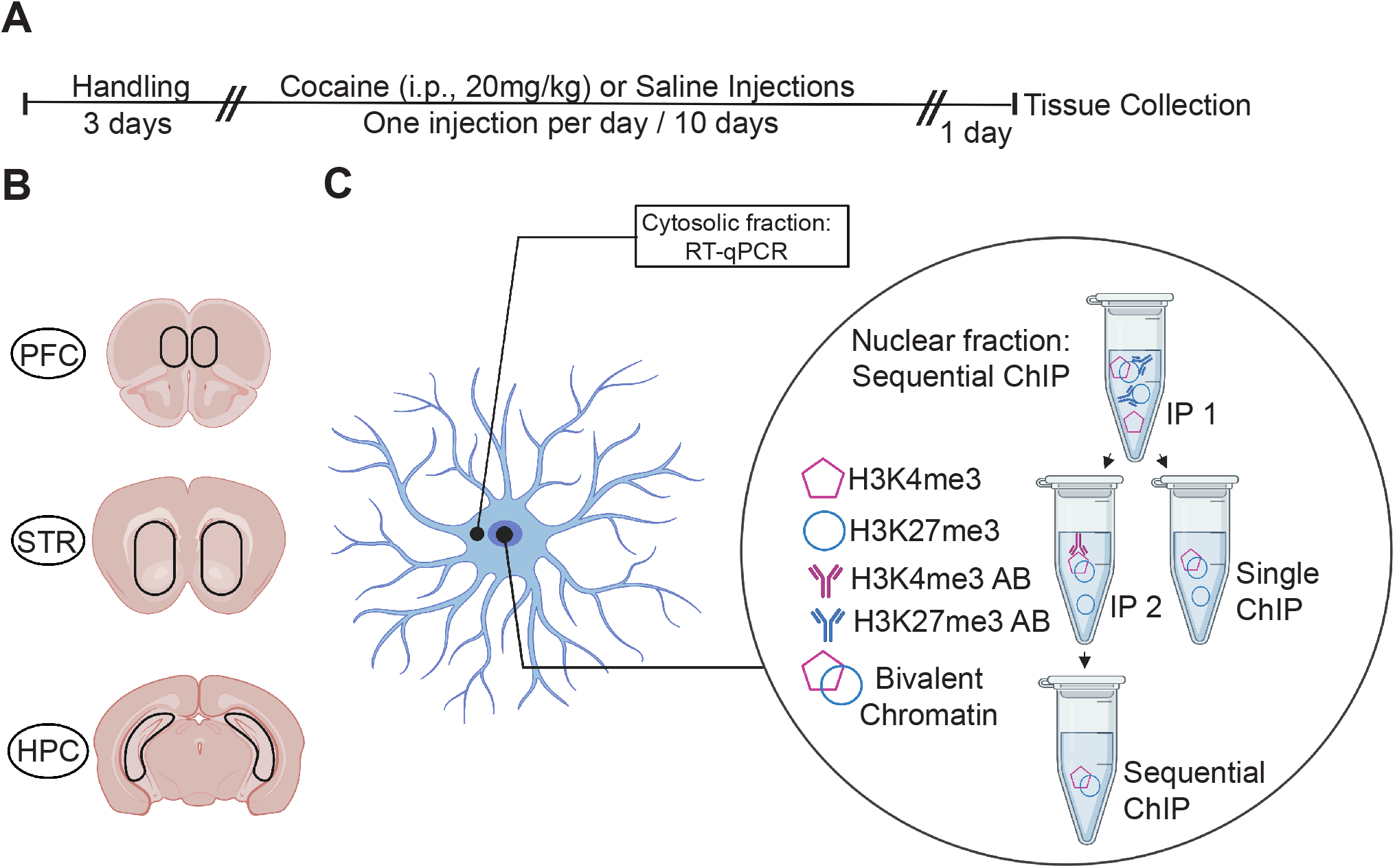
Methods used to investigate regional transcriptional and epigenomic profiles following cocaine exposure. **(A)** Timeline for investigator-administered cocaine (20 mg/kg; i.p) paradigm. **(B)** Coronal brain slice illustrations showing the brain regions taken for single sample sequencing (Top to bottom: prefrontal cortex (PFC), striatum (STR), hippocampus (HPC)). **(C)** Schematic showing the sequential ChIP protocol that includes two immunoprecipitations.

### S3EQ nuclear and cytosolic fraction separation

S3EQ (33) was conducted as previously published with modified buffer volume. Tissue samples were homogenized in 350 μL cell lysis buffer (10 mM Tris-HCl (pH 8.0), 10 mM NaCl, 3 mM MgCl2 and 0.5% NP-40 in H2O) and spun for five minutes (1,500 x g, 4°C). The nuclei-containing pellet and RNA-containing cytosolic supernatant were separated and subjected to sequential ChIP or RNA extraction, respectively **(Figure 1C)**.

### Single and Sequential Chromatin Immunoprecipitation (ChIP) and ChIP-PCR (qCHIP)

Prior to initiating the protocol, the following solutions were prepared as previously described (33,34): <u>BSA Blocking Buffer</u> (0.5% Bovine serum albumin in 1x PBS), <u>Dilution Buffer</u> (16.7 mM Tris-HCl (pH 8.0), 1.1% Triton-X 100, 0.01% SDS, 167 mM NaCl, 1.2mM EDTA in H_2_O), <u>Nuclear Lysis Buffer</u> (50 mM Tris-HCl (pH 8.0), 5 mM EDTA, 1% SDS in H_2_O), <u>Wash Buffer 1</u> (20 mM Tris-HCl pH 8.0, 150 mM NaCl, 2 mM EDTA, 1% Triton X-100, 0.1% SDS in H_2_O), <u>Wash Buffer 2</u> (20 mM Tris-Cl pH 8.0, 500 mM NaCl, 2 mM EDTA, 1% Triton X-100, 0.1% SDS in H_2_O), <u>Wash Buffer 3</u> (250 mM LiCl, 10 mM Tris-HCl pH 8.0, 1% sodium deoxycholate, 1 mM EDTA, 1% IGEPAL CA-630 in H_2_O), <u>TE Buffer</u> (10 mM Tris-HCl pH 8.0, 1 mM EDTA in H_2_O), <u>Elution Buffer</u> (0.1 M NaHCO_3_, 1% SDS in H_2_O).

M280 Sheep anti-Rabbit Dynabeads (Invitrogen, 11204D) were washed three times in BSA blocking buffer. The beads were bound to either H3K4me3 antibody (Rabbit Polyclonal, EMD Millipore, 07-473) or H3K27me3 antibody (Rabbit Polyclonal, EMD Millipore, 07-449) in 2X bead volume of dilution buffer and rotated for six hours (4°C). Nuclei obtained from S3EQ were resuspended in PBS and fixed with 1% formaldehyde for eight minutes (350 rpm, 22°C). 100 μL 1 M glycine was added and samples were rocked for five minutes to stop cross-linking (500 rpm, 22°C). Samples were then centrifuged for five minutes (5500 x g, 4°C) and the supernatant was discarded. The pellet was resuspended in 200 μL nuclear lysis buffer, transferred to TPX tubes, (Diagenode, C30010010) and incubated on ice for ten minutes. Samples were then sonicated in a Bioruptor® (Diagenode) for two runs (high setting, 30s on, 30s off, 15 cycles). Dilution buffer was added to reach 1000 μL. 10% input (relative to each IP) was collected from each sample for normalization later used in qChIP analysis. The remaining diluted chromatin was combined with antibody-bound beads (1.2 μL bead-antibody slurry/1 μL chromatin), dilution buffer, and placed on a rotator (O/N, 4°C). Samples were then washed with 1 mL of ice-cold Wash Buffer 1, Wash Buffer 2, Wash Buffer 3, and TE Buffer. Samples were rotated for five minutes during each wash (RT). Following washes, samples were rocked with 200 μL of elution buffer for 20 minutes (500 rpm, RT), centrifuged for three minutes (14,000 x g, RT), and placed on a magnet. The supernatant from each sample was divided in half and transferred to fresh tubes, each containing 100 μL of elution buffer. One tube was collected as the single ChIP sample.

The remaining tube was subjected to a second immunoprecipitation (sequential ChIP samples), in which each sample was subjected to antibody-bound beads that were bound to the reciprocal antibody (i.e. H3K4me3 ChIP samples were bound to H3K27me3 antibody-bound beads) and placed on a rotator (O/N, 4°C). The sequential ChIP samples were processed according to the single ChIP protocol above. Single and sequential ChIP samples were then incubated for 4 hours (65°C) following the addition of 8 μL 5 M NaCl. Proteinase K (0.002 mg) was added, and incubation continued for two hours (65°C). Samples were heat inactivated (15 minutes, 78°C) and DNA was purified using the QiaAmp DNA Micro kit (Qiagen, 56304). qChIP was conducted as previously published (26). Briefly, qChIP was run using Power SYBR™ Green PCR Master Mix (Life Technologies). The primer sequences that span −100 to +100bps of target promoter regions used to assess the presence of HPTMs (26) are noted in **Supplemental Table 1**.

### RNA Extraction and qPCR

The cytosolic fraction following S3EQ was mixed with 600 μL RLT buffer (Qiagen) and RNA was extracted using the RNAeasy Micro Kit (Qiagen, 74004). RNA concentration was obtained using a Qubit 4 Fluorometer and RNA HS Assay Kit (Invitrogen). cDNA was synthesized using the Bio-Rad iScript™ cDNA Synthesis Kit (Bio-Rad, 1708890). qPCR was conducted using Power SYBR™ Green PCR Master Mix (Life Technologies). The primer sequences used to assess mRNA levels are noted in **Supplemental Table 2**.

### Pearson Correlation analysis of qCHIP and qPCR

Pearson correlation analysis was used to assess the relationship between K4&K27 bivalent *Nr4a1* promoter, *Nr4a1* or *Cartpt* mRNA levels, and *Nr4a1* or *Cartpt* monovalent H3K4me3 or H3K27me3. Scripts were written and executed in R v4.1.1. First, all values corresponding to relevant endpoints were compiled for each sex (male, female) and brain region (STR, HPC, PFC) combination. For a given combination, the dataset was further separated based on treatment (saline, cocaine). Analyses were conducted with input values (fold change) from qChIP and qPCR data. An endpoint (qChIP or qPCR) was excluded if it lacked at least three observations in both the saline and cocaine datasets. An all-against-all Pearson’s correlation matrix was generated for the saline and cocaine datasets separately using the base R cor(method=”pearson”) function. A correlation heatmap was generated from the Pearson correlation matrix using the corrplot() function from the corrplot package.

### Statistical Analyses

Statistical tests used reflect the experimental design. Specifically, while male and female mice were treated with cocaine in a single cohort, the tissue for each sex and each brain region was processed and analyzed separately due to the technical limitations associated with processing large numbers of sample. We therefore did not apply ANOVA to analyze sex or region differences. The Grubbs test was used to identify all outliers. Student’s t-tests were used for qChIP and qPCR analyses as these experiments directly compared one factor (drug exposure) from two groups. Sex was not considered a variable within any analyses due to experimental execution. F-tests of variance were conducted in every analysis and Welch’s correction was utilized as needed. These data are shown as mean ±SEM. Graphpad Prism V9.3 was used for qChIP and qPCR analyses. qChIP data was analyzed by comparing CT values of each experimental group versus the control group following normalization of each sample’s input using the **ΔΔ**Ct method (35,36). Bivalency at *Cartpt* was not assessed, as not all sequential-immunoprecipitated samples reached minimum signal intensity for qChIP. qPCR data was analyzed by comparing CT values of each experimental group versus the control group following normalization of a housekeeping gene (*Gapdh*) using the **ΔΔ**Ct method (35,36). All experiments were carried out one to two times, and data replication was observed in all instances of repeated experiments. Additionally, every qChIP and qPCR dataset had a technical replicate conducted by an independent investigator to ensure the validity of the data. Regarding visual data representation, each data point in all graphs represent one animal **(Figures 2-5)**. For Pearson correlation analysis, the p-value matrix of all correlations corresponding to each comparison was extracted using the two-sided cor.mtest(method=”pearson”) function from the corrplot package with a 95% confidence interval. This statistical test for Pearson’s product-moment coefficient was calculated following a t-distribution (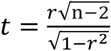, where n = number of observations and r = calculated Pearson correlation coefficient) with degrees of freedom equal to length (number of complete observations) – 2. Heatmap tiles were shaded red or blue only when satisfying a p-value ≤ 0.05 using p.mat=p.mat_for_comparison and sig.level=0.05 within the corrplot() function. Heatmap tiles were shaded gray when analysis was not conducted, as analysis would violate the rule of independence (37). All scripts are available at: (https://github.com/HellerLAbeats/Nr4a1_bivalency)

**Figure 2.**
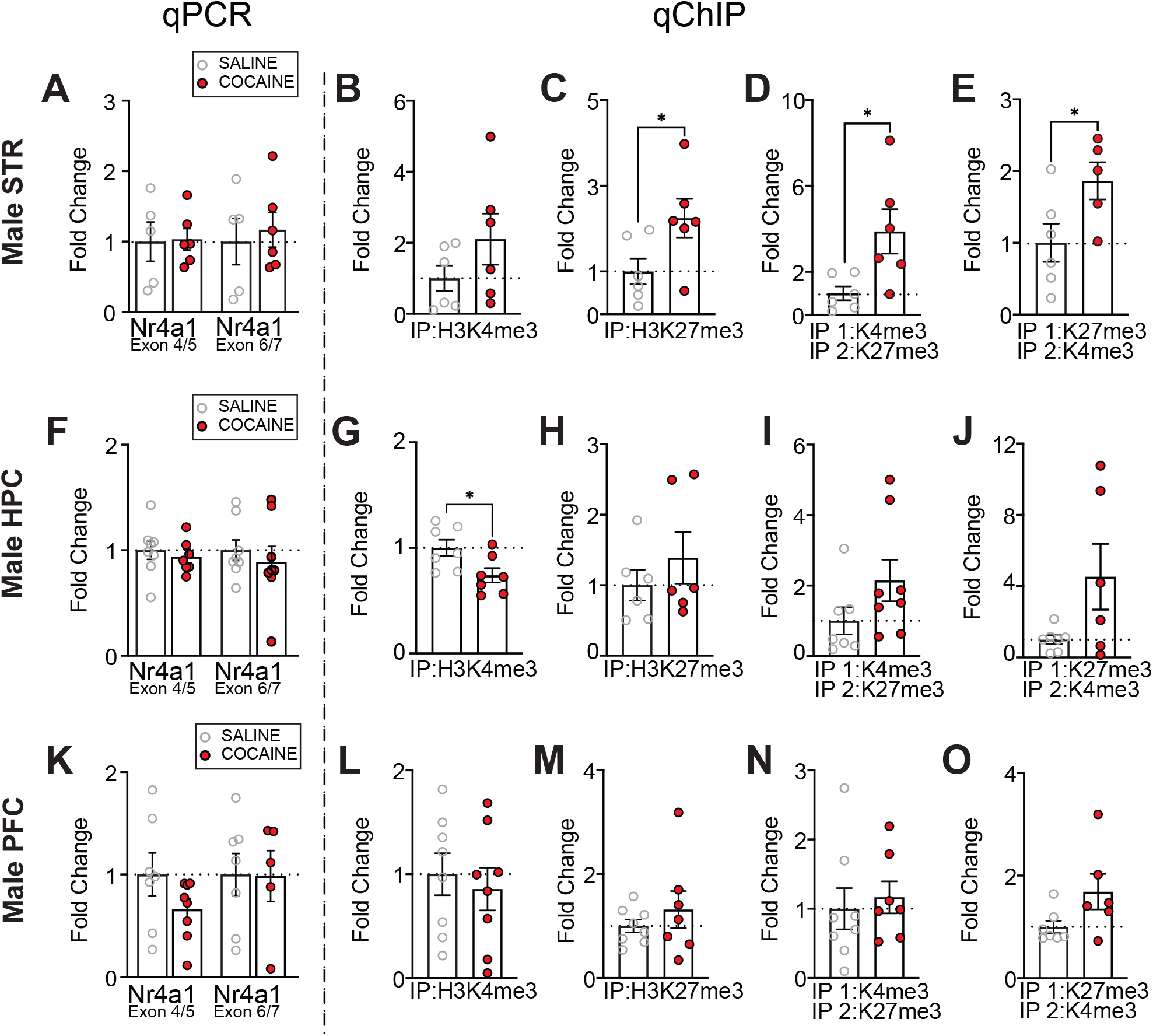
Cocaine alters the epigenetic profile of *Nr4a1* in the striatum and hippocampus of male mice. **(A)** No change in mRNA levels of *Nr4a1* (as examined by primer pairs that cover the junction between exon 4/5 and exon 6/7) were observed in the male mouse STR following 10 days of investigator-administered cocaine (n = 5-6/group). There was no change in **(B)** H3K4me3 (n = 6/group), but an increase in **(C)** H3K27me3 (n = 6/group), **(D)** H3K4me3/H3K27me3 bivalency (n = 6/group), and **(E)** H3K27me3/H3K4me3 bivalency (n = 5-6/group) at *Nr4a1* in the male mouse STR following 10 days of investigator-administered cocaine. **(F)** There was no change in mRNA levels of *Nr4a1* (as measured by independent primer pairs that cover the junction between exons 4/5 and exons 6/7) in the male mouse HPC following 10 days of investigator-administered cocaine (n = 7-8/group). There was a decrease in **(G)** H3K4me3 at (n = 7/group), but no change in **(H)** H3K27me3 (n = 6/group), **(I)** H3K4me3/H3K27me3 bivalency (n = 7-8/group), or **(J)** H3K27me3/H3K4me3 bivalency (n = 6-7/group) at *Nr4a1* in the male mouse HPC following 10 days of investigator-administered cocaine. **(K)** There was no change in mRNA levels of *Nr4a1* (as measured by independent primer pairs that cover the junction between exons 4/5 and exons 6/7) in the male mouse PFC following 10 days of investigator-administered cocaine (n = 7-8/group). There is no change in **(L)** H3K4me3 (n = 8/group), **(M)** H3K27me3 (n = 7-8/group), **(N)** H3K4me3/H3K27me3 bivalency (n = 7-8/group), or **(O)** H3K27me3/H3K4me3 bivalency at *Nr4a1* in the male mouse PFC following 10 days of investigator-administered cocaine (n = 6-7/group). *p < 0.05. Data are shown as mean ± SEM. STR, striatum; HPC, hippocampus; PFC, prefrontal cortex.

## RESULTS

### Sequential ChIP-qPCR demonstrated K4&K27 bivalency at *Nr4a1* promoter in brain reward regions is present, in both male and female mice

For epigenetic profiling of three reward-associated brain regions following cocaine treatment, we applied sequential ChIP to examine K4&K27 bivalency at *Nr4a1* **(Figure 1)**. ChIP-qPCR and ChIP-Seq protocols that quantify HPTM enrichment of monovalent promoters are unable to quantify bona fide combinatorial HPTM enrichment at single promoter. This weakness stems from the fact that individual HPTM profiling experiments cannot distinguish between bivalent promoter enrichment of two HPTMs in a single population of cells or monovalent promoter enrichment of two HPTMs from a mixed population of cells (see **Figure 2** in (38)). The sequential ChIP protocol is required to define bona fide bivalency as H3K4me3 and H3K27me3 enrichment at the same locus in a single population of cells (and nucleosomes) (38). To measure bivalency, we split each single sample and performed H3K4me3 ChIP followed by H3K27me3 ChIP. From the same sample, we also performed H3K27me3 ChIP followed by H3K4me3 ChIP. In this way, sequential ChIP was assayed with either H3K4me3 or H3K27me3 as the first IP, and the other HPTM as the second IP. This approach allowed us to interpret differences in cocaine-regulated K4&K27 bivalency due to differences in rates of deposition or removal between the two HPTMs (39). In each case, enrichment based on antibody order was quantified by qCHIP. As we found no differences in results of qCHIP by order of sequential ChIP, we concluded that both approaches demonstrated *Nr4a1* K4&K27 promoter bivalency.

### Cocaine enriched H3K27me3 and bivalency at the *Nr4a1* promoter and did not alter *Nr4a1* mRNA levels in male striatum

Following cocaine treatment, there was no change in *Nr4a1* mRNA **(Figure 2A)** or H3K4me3 enrichment at *Nr4a1* in the male STR **(Figure 2B)**. In the same samples there was enrichment of H3K27me3 at *Nr4a1* **(Figure 2C)** (t_10_ = 2.3, p = 0.0443) and an increase in K4&K27 bivalency at the *Nr4a1* promoter **(Figure 2D, E)** (**D**: t_5.973_ = 2.663, p = 0.0376; **E**: t_9_ = 2.298, p = 0.0476). In the male HPC following cocaine treatment, there was no change in *Nr4a1* mRNA **(Figure 2F)**, but we observed deenrichment of H3K4me3 at *Nr4a1* **(Figure 2G)** (t_12_ = 2.528, p = 0.0265). In the same samples, there was no change in H3K27me3 enrichment or K4&K27 bivalency **(Figure 2H, I, J)**. In the male PFC following cocaine treatment, there was no change in *Nr4a1* mRNA **(Figure 2K)**, H3K4me3, H3K27me3, or K4&K27 bivalency **(Figure 2L-O)**.

### Cocaine increased *Cartpt* mRNA specifically in PFC of male mice

Given the transient nature of immediate early gene expression, we also measured mRNA and HPTM enrichment of a downstream transcriptional target of *Nr4a1* (26). We found no changes in *Cartpt* mRNA, or that of other dopamine-related targets - dopamine receptors, *Drd1* and *Drd2*, or adenosine A2a receptor (*A2a*), in the male STR following cocaine treatment **(Figure 3A)**. Likewise, we observed no change in H3K4me3 or H3K27me3 enrichment at *Cartpt* **(Figure 3B, C)**. Similarly, in the male HPC, cocaine treatment did not alter *Cartpt, Drd1, Drd2*, or *A2a* mRNA **(Figure 3D)**, or H3K4me3 or H3K27me3 enrichment at *Cartpt* **(Figure 3E, F)**. Alternatively, in the male PFC, cocaine treatment increased *Cartpt* mRNA (t_7.165_ = 3.182, p = 0.0150), and we observed a trending decrease of *Drd1* mRNA (trending, t_7.247_ = 2.059, p = 0.0771), *Drd2* (trending, t_7.318_ = 2.110, p = 0.0711), but cocaine did not alter *A2a* mRNA levels **(Figure 3G)**. This was accompanied by no change in H3K4me3 or H3K27me3 enrichment at *Cartpt* **(Figure 3H, I)**. These data suggest cocaine induces region-specific expression of *Cartpt* in the male PFC, but not in STR or HPC.

**Figure 3.**
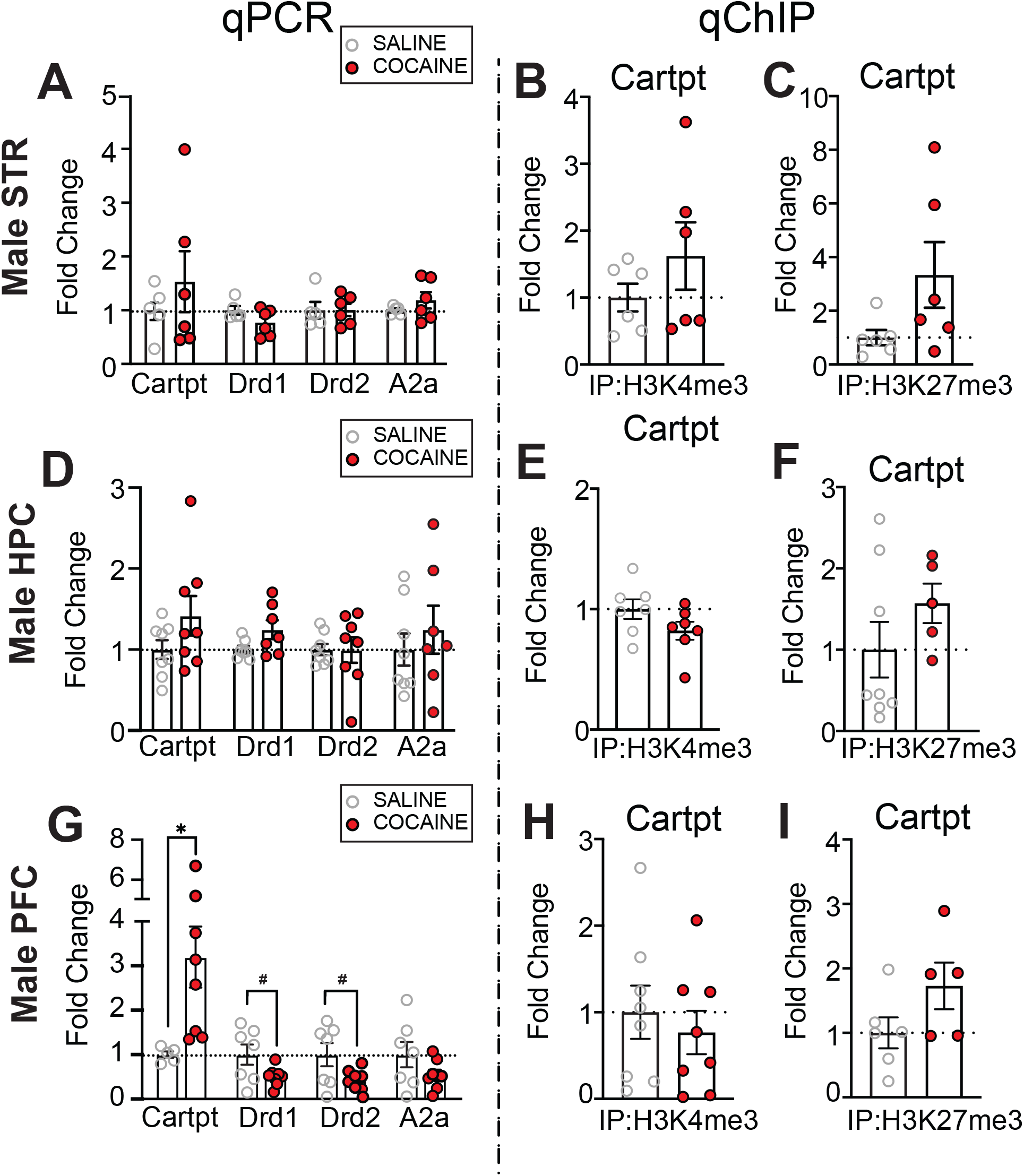
*Nr4a1* downstream target, *Cartpt*, is regionally activated after 10 days of investigator-administered cocaine in male mice. **(A)** There was no change in mRNA levels of *Cartpt, Drd1, Drd2*, or *A2a* in the male mouse STR following 10 days of investigator-administered cocaine (n = 5-6/group). **(B)** There was no change in **(B)** H3K4me3 (n = 6/group) or (**C)** H3K27me3 (n = 6/group) at *Cartpt* in the male mouse STR after 10 days of investigator-administered cocaine. **(D)** There was no change in mRNA levels of *Cartpt, Drd1, Drd2*, or *A2a* in the male mouse HPC following 10 days of investigator-administered cocaine (n = 7-8/group). **(E)** There was no change in **(E)** H3K4me3 (n = 7/group) or **(F)** H3K27me3 (n = 6-8/group) at *Cartpt* in the male mouse HPC after 10 days of investigator-administered cocaine. **(G)** There was an increase in mRNA levels of *Cartpt*, a trending decrease in mRNA levels of *Drd1* and *Drd2*, and no change in mRNA levels of *A2a* in the male mouse PFC following 10 days of investigator-administered cocaine (n = 5-6/group). There was no change in **(H)** H3K4me3 (n = 8/group) or **(I)** H3K27me3 at *Cartpt* in the male mouse PFC after 10 days of investigator-administered cocaine (n = 5-6/group). *p < 0.05. ^#^p < 0.10. Data are shown as mean ± SEM. STR, striatum; HPC, hippocampus; PFC, prefrontal cortex.

### Cocaine increased *Nr4a1* mRNA in multiple reward-associated brain regions in female mice

Following cocaine treatment, *Nr4a1* mRNA was increased in the female STR **(Figure 4A)**. This increase was independently identified using two different primer sets, each of which spans the junction between two distinct exons within *Nr4a1* (*Nr4a1* Exon 4/5: t_9_ = 2.771, p = 0.0217; *Nr4a1* Exon 6/7: t_9_ = 3.030, p = 0.0142). In the same samples, there was no change in H3K4me3, H3K27me3, or K4&K27 bivalency at *Nr4a1* **(Figure 4B-E)**. In the female HPC, there was no change in *Nr4a1* mRNA following cocaine treatment when the *Nr4a1* Exon 4/5 primer set was used, but we observed a trending increase in *Nr4a1* mRNA when mRNA levels were evaluated with Exon 6/7 primers (trending, t_14_ = 1.1841, p = 0.0869) **(Figure 4F)**. In the same samples, there was no change in *Nr4a1* H3K4me3, H3K27me3 or K4&K27 bivalency **(Figure 4G-J)**. In the female PFC following cocaine treatment, *Nr4a1* mRNA was increased when measured at *Nr4a1* E6/7 but not *Nr4a1* E4/5 (*Nr4a1* Exon 6/7: t_13_ = 2.327, p = 0.0368) **(Figure 4K)**. In the same samples, there was no change in H3K4me3 **(Figure 4L)**, there was H3K27me3 enrichment (t_6.392_ = 3.363, p = 0.0138) **(Figure 4M)**, and no change in K4&K27 bivalency **(Figure 4N, O)**. In sum, these data showed that in male STR cocaine treatment did not alter *Nr4a1* mRNA but did increase H3K27me3 and K4&K27 bivalency at the *Nr4a1* promoter. In contrast, cocaine treatment in female mice increased *Nr4a1* mRNA expression in the STR along with maintenance of H3K4me3, H3K27me3, and K4&K27 bivalency at the *Nr4a1* promoter.

**Figure 4.**
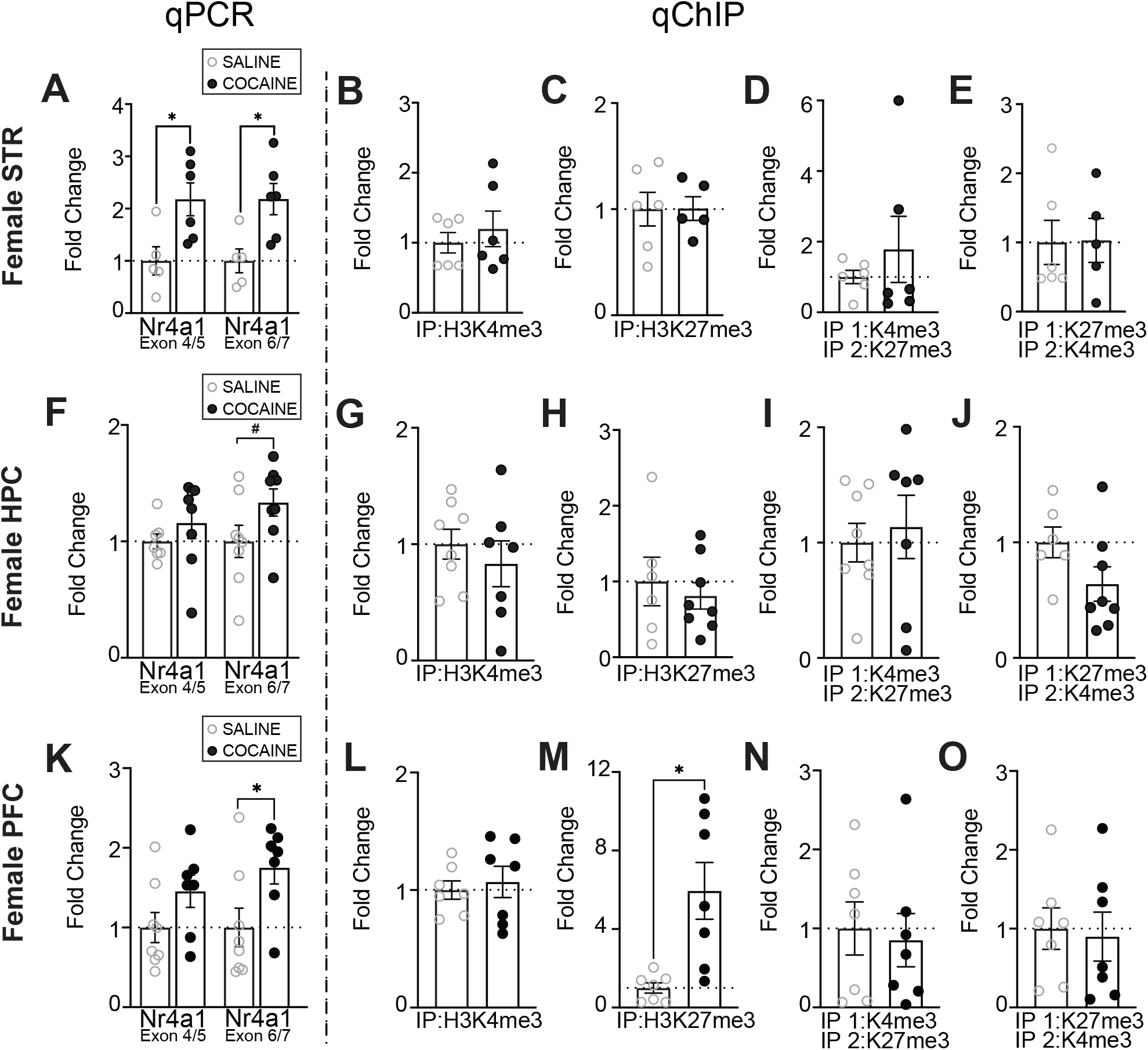
Cocaine alters the transcriptomic profile in the STR, and the transcriptomic and epigenetic profile of *Nr4a1* in the PFC of females. **(A)** There was an increase in mRNA levels of *Nr4a1* (as examined by primer pairs that cover the junction between exons 4/5 and exons 6/7) in the female mouse STR following 10 days of investigator-administered cocaine (n = 4-6/group). There was no change in **(B)** H3K4me3 (n = 6/group), **(C)** H3K427me3 (n = 5-6/group), **(D)** H3K4me3/H3K27me3 bivalency (n = 6/group), or **(E)** H3K27me3/H3K4me3 bivalency (n = 5-6/group) at *Nr4a1* in the female mouse STR after 10 days of investigator-administered cocaine. **(F)** There was no change in mRNA levels of *Nr4a1* as measured by independent primer pairs that cover the junction between exons 4/5, and a trending increase in mRNA levels of *Nr4a1* as measured by independent primer pairs that cover the junction between exons 6/7 in the female mouse HPC following 10 days of investigator-administered cocaine (n = 7-8/group). There was no change in **(G)** H3K4me3 (n = 7-8/group), **(H)** H3K27me3 (n = 6-8/group), **(I)** H3K4me3/H3K27me3 (n = 7-8/group), or **(J)** H3K27me3/H3K4me3 bivalency (n = 6-8/group) at *Nr4a1* in the female mouse HPC following 10 days of investigator-administered cocaine. **(K)** There was an increase in mRNA levels of *Nr4a1* (as measured by independent primer pairs that cover the junction between exons 6/7, but not exons 4/5) in the female mouse PFC following 10 days of investigator-administered cocaine (n = 7-8/group). There was no change in **(L)** H3K4me3 (n = 7/group), but there was an increase in **(M)** H3K27me3 (n = 7/group) at *Nr4a1* following 10 days of investigator-administered cocaine in the female mouse PFC (n = 7/group). **(N)** There was no change in H3K4me3/H3K27me3 bivalency (n = 7/group) or **(O)** H3K27me3/H3K4me3 bivalency at *Nr4a1* in the female mouse prefrontal cortex following 10 days of investigator-administered cocaine (n = 7/group). *p < 0.05. ^#^p < 0.10. Data are shown as mean ± SEM. STR, striatum; HPC, hippocampus; PFC, prefrontal cortex.

### Cocaine did not alter mRNA in drug-related markers, *Cartpt* mRNA, or *Cartpt* HPTMs in female mouse brain regions

As we saw multiple regional changes in mRNA levels of *Nr4a1* following cocaine in female mice, we expected to find transcriptomic or epigenetic changes at *Cartpt*, a downstream target of *Nr4a1* (26). Surprisingly, we found no transcriptomic or epigenetic changes (H3K4me3, H3K27me3) at *Cartpt* in female mice following cocaine in the STR **(Figure 5A-C)**, HPC **(Figure 5D-F)**, or PFC **(Figure 5G-I)**, although we observed trending deenrichment of H3K27me3 levels at *Cartpt* in the PFC **(Figure 5I)** (t_11_ = 1.953, p = 0.0767). Notably, we also observed no change in mRNA levels of *Drd1, Drd2*, or *A2a* in the STR, HPC, or PFC **(Figure 5A, D, G)** in female mice following cocaine, but observed a trending decrease in mRNA levels of *Drd1* in the PFC **(Figure 5G)** (t_13_ = 1.887, p = 0.0818).

**Figure 5.**
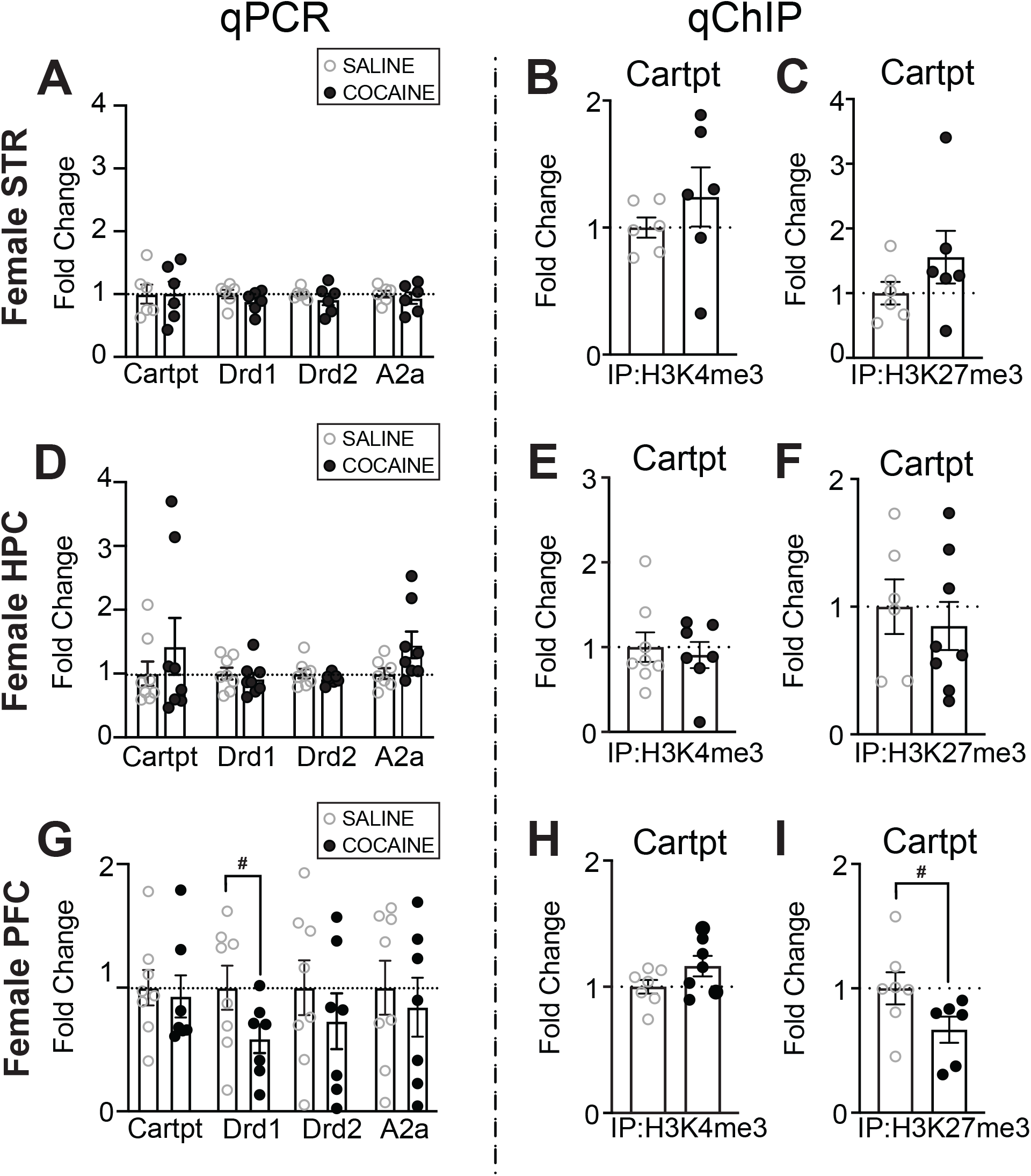
*Nr4a1* downstream target, *Cartpt*, and other drug-related markers are selectively impacted after 10 days of investigator-administered cocaine in female mice. **(A)** There was no change in mRNA levels of *Cartpt, Drd1, Drd2*, or *A2a* in the female mouse STR following 10 days of investigator-administered cocaine (n = 6/group). There was no change in **(B)** H3K4me3 (n =6/group) or **(C)** H3K27me3 (n = 6/group) at *Cartpt* in the female mouse STR after 10 days of investigator-administered cocaine. **(D)** There was no change in mRNA levels of *Cartpt, Drd1, Drd2*, or *A2a* in the female mouse HPC following 10 days of investigator-administered cocaine (n = 8/group). There was no change in **(E)** H3K4me3 (n = 7-8/group) or **(F)** H3K27me3 (n = 6-8/group) at *Cartpt* in the female mouse HPC after 10 days of investigator-administered cocaine. **(G)** There was a trending decrease in mRNA levels of *Drd1* and no change in mRNA levels of *Cartpt, Drd2*, or *A2a* in the female mouse PFC following 10 days of investigator-administered cocaine (n = 7-8/group). There was no change in **(H)** H3K4me3 (n = 7-8/group) but there was a trending decrease in **(I)** H3K27me3 at *Cartpt* in the female mouse PFC after 10 days of investigator-administered cocaine (n = 7-8/group). *p < 0.05. ^#^p < 0.10. Data are shown as mean ± SEM. STR, striatum; HPC, hippocampus; PFC, prefrontal cortex.

### Cocaine impacts correlational relationships within and between the transcriptomic and epigenetic profiles of *Nr4a1* and *Cartpt* in male and female mice

As we hypothesized that we would see a linear relationship between *Nr4a1* and *Cartpt* following cocaine, we utilized Pearson’s correlation matrices to explore the relationships within our sample population between the transcriptomic and epigenetic profiles of *Nr4a1* and *Cartpt*. In the STR we observed positive correlations between (1) *Nr4a1* H3K4me3 and *Cartpt* H3K4me3 and (2) mRNA levels of *Nr4a1* measured by the two primers pairs spanning either Exon 4/5 or Exon 6/7 across treatments (saline, cocaine) and sex (male, female) **(Figure 6A, B)**. In the STR of saline-treated females, we observed a positive correlation between (3) *Nr4a1* mRNA and *Cartpt* H3K27me3 **(Figure 6B)**. In the STR of males following cocaine, we observed a positive correlation between (4) *Cartpt* mRNA and *Cartpt* H3K27me3, (5) *Cartpt* mRNA and *Nr4a1* K4/K27 bivalency, and (6) *Cartpt* H3K27me3 and *Nr4a1* K4/K27 bivalency **(Figure 6A)**. In the STR of females following cocaine, we observed a positive correlation between (7) *Nr4a1* H3K4me3 and H3K4me3 at *Cartpt* and (8) *Nr4a1* H3K27me3 and *Cartpt* H3K4me3 **(Figure 6B)**.

**Figure 6.**
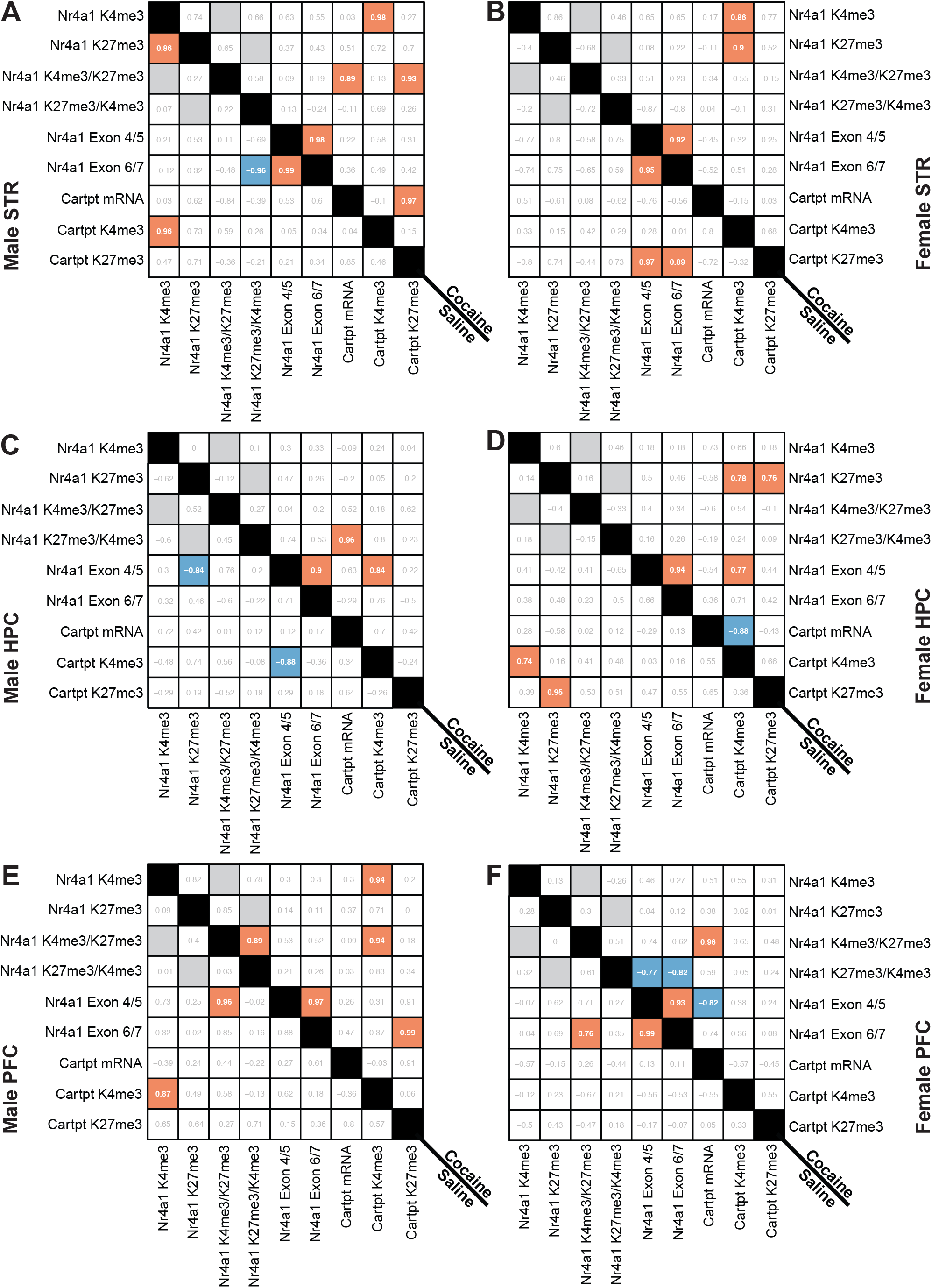
Pearson’s Correlation Matrices reveal distinct transcriptomic and epigenetic relationships following cocaine exposure in both sexes. **(A)** Pearson’s Correlation matrix evaluating transcriptomic and epigenetic relationships of *Nr4a1* and *Cartpt* in the striatum of saline- and cocaine-injected males and **(B)** females. **(C)** Pearson’s Correlation matrix evaluating transcriptomic and epigenetic relationships of *Nr4a1* and *Cartpt* in the hippocampus of saline- and cocaine-injected males and **(D)** females. **(E)** Pearson’s Correlation matrix evaluating transcriptomic and epigenetic relationships of *Nr4a1* and *Cartpt* in the prefrontal cortex of saline- and cocaine-injected males and **(F)** females. For all Pearson’s Correlation matrices, red indicates a significant positive correlation (p ≤ 0.05) and blue indicates a significant negative correlation (p ≤ 0.05). A gray box indicates a correlation was not analyzed as it validates the rule of independence for Pearson’s Correlation.

In the HPC of saline-treated male mice we observed a negative correlation between (1) *Nr4a1* mRNA levels (Exon 4/5) and *Nr4a1* H3K27me3 and (2) *Nr4a1* mRNA levels (Exon 4/5) and *Cartpt* H3K4me3 **(Figure 6C)**. In the HPC of saline-treated females we observed a positive correlation between (3) *Nr4a1* H3K4me3 and *Cartpt* H3K4me3 and (4) *Nr4a1* H3K27me3 and *Cartpt* H3K27me3 **(Figure 6D)**. In the HPC of males following cocaine, we observed a positive correlation between (5) *Nr4a1* mRNA levels (Exon 4/5 and Exon 6/7), (6) *Nr4a1* mRNA levels (Exon 4/5) and *Cartp*t H3K4me3, and (7) *Cartpt* mRNA levels and *Nr4a1* K27/K4 bivalency **(Figure 6C)**. In the HPC of females following cocaine, we observed a positive correlation between (8) *Nr4a1* mRNA levels (Exon 4/5 and Exon 6/7), (9) *Nr4a1* mRNA levels (Exon 4/5) and *Cartp*t H3K4me3, (10) *Nr4a1* H3K27me3 and *Cartpt* H3K4me3, and (11) *Nr4a1* H3K27me3 and *Cartpt* H3K27me3 **(Figure 6D)**. We observed a negative correlation in the HPC of females following cocaine between (12) *Cartpt* mRNA levels and *Cartpt* H3K4me3 **(Figure 6D)**.

In the PFC of saline-treated males, we found a positive correlation between (1) mRNA levels of *Nr4a1* (Exon 4/5) and H3K4me3/H3K27me3 at *Nr4a1* and (2) H3K4me3 at *Nr4a1* and H3K4me3 at *Cartpt* **(Figure 6E)**. In the PFC of saline-treated females, we found a positive correlation between (3) *Nr4a1* mRNA levels (Exon 4/5 and Exon 6/7) as well as (4) *Nr4a1* mRNA levels (Exon 6/7) and *Nr4a1* K4/K27 bivalency **(Figure 6F)**. In the PFC of cocaine-injected males, we found multiple positive correlations including: (5) H3K4me3/H3K27me3 bivalency at *Nr4a1* and H3K4me3 at *Cartpt*, (6) H3K4me3 at *Nr4a1* and H3K4me3 at *Cartpt*, (7) *Nr4a1* H3K4me3/H3K27me3 bivalency, (8) *Nr4a1* mRNA levels (Exon 4/5 and Exon 6/7), and (9) *Nr4a1* mRNA levels (Exon 6/7) and *Cartpt* H3K27me3 **(Figure 6E)**. In the PFC of females following cocaine, we observed both positive and negative correlations. We found that (10, 11) mRNA levels of *Nr4a1* (Exon 4/5 and Exon 6/7) negatively correlate with H3K27me3/H3K4me3 bivalency at *Nr4a1*. Additionally, (12) mRNA levels of *Nr4a1* (Exon 4/5) negatively correlate with mRNA levels of *Cartpt* **(Figure 6F)**. We observed a positive correlation between (13) mRNA levels of *Cartpt* and H3K4me3/H3K27me3 bivalency at *Nr4a1* and (14) mRNA levels of *Nr4a1* (Exon 4/5 and Exon 6/7) **(Figure 6F)**.

**Supplementary Tables 3-5** contain every significant correlation observed, and the corresponding figure(s) that show the data within each correlation in **Figure 6** (ex. the data used to reveal the positive correlation between *Nr4a1* mRNA and *Cartpt* H3K27me3 in the STR of saline-treated females can been seen in **Figure 4A and Figure 5C**.) Importantly, the relationships with bivalency (H3K4me3/H3K27me3 vs H3K27me3/H3K4me3) do not always align at the level of a correlation. This is likely due to the biological mechanism of bivalency. Induction of H3K4me3/H3K27me3 bivalency is indicative of a transition from an activating to poised state (bivalency results in lower activation), while induction of H3K27me3/H3K4me3 bivalency is indicative of a repressive to poised state (bivalency results in higher activation).

## DISCUSSION

There is an emerging literature on the transcriptomic and epigenetic profiles of *Nr4a1* and its regulation by cocaine. Despite this progress, no study to date has explored homeostatic and cocaine-activated regulation of *Nr4a1* in both males and females in any organism. Here, we demonstrate that, following cocaine, *Nr4a1* mRNA increases in female mouse STR and PFC, and does not change in male mouse STR, HPC, or PFC. We also find an increase in *Nr4a1* K4&K27 promoter bivalency in the male STR following cocaine treatment. Finally, we show that the mRNA level of a putative *Nr4a1* target, *Cartpt*, is activated specifically in the male PFC.

The current study profiles both ventral and dorsal STR, as well as HPC and PFC, other brain regions associated with reward neuropathology. Both the ventral STR (NAc) and dorsal STR are involved in cocaine reinforcement, and the interconnectivity between these regions mediates cocaine-seeking behavior (40). Our lab and others find that cocaine treatment increases *Nr4a1* mRNA in the NAc in mixed sex populations (26), as well as in males and females when assessed separately (27). However, in order to analyze K4&K27 bivalency at the *Nr4a1* promoter of a single mouse, it was necessary to isolate chromatin from both ventral and dorsal STR and perform sequential ChIP. New methods are being developed to allow sequential ChIP profiling of small amounts of input material and to allow profiling of more specific brain regions and cell-types. For example, recently developed multi-CUT&Tag (41) allows for mapping of HPTM co-localization in the same cell, and single-cell multi-CUT&Tag (41) can probe for bivalency in specific cell types. While single-cell CUT&Tag has been carried out in mouse brain to profile individual HPTMs (42), these multi-CUT&Tag methods that uncover bivalency have yet to be applied in brain tissue to the best of our knowledge.

To date, few studies have assessed bivalency in the brain (23–25), and none, to our knowledge, have investigated cocaine regulation of bivalency. Bivalent chromatin regulates DNA accessibility rather elegantly by transitioning euchromatin or heterochromatin to a poised, bivalent state that can quickly react to cellular cues. This flexibility is beneficial for cell differentiation (22) and stress responses (23), but may underlie detrimental instability of gene expression in the adult brain. Consequently, bivalency is implicated in transcriptional dysregulation observed in Huntington’s Disease (43) as well as multiple types of cancer (44–47). We hypothesize that bivalency may similarly underlie transcriptional dysregulation in drug addiction, in both males and females. Our data highlight that the interplay between bivalency and cocaine warrants further investigation.

Interestingly, we found that *Nr4a1* mRNA levels did not always correspond with the canonical roles of H3K4me3, H3K27me3, and K4&K27 bivalency, in activating, repressing, and poising gene expression, respectively. For example, in the female PFC following cocaine treatment we observed concomitant increases in *Nr4a1* mRNA and H3K27me3 at *Nr4a1*. We thus applied correlational analyses to examine these relationships more comprehensively. In the male STR, we found a positive correlation between *Nr4a1* K4/K27 bivalency and Cartpt mRNA levels. Given that *Nr4a1* is a putative activator of *Cartpt* expression (26), this finding went against our expectation that *Nr4a1* gene poising would be negatively correlated with *Cartpt* mRNA levels. Additionally, given the lack of transcriptomic or the epigenomic regulation of *Nr4a1* in the male PFC and the vast increase in *Cartpt* mRNA levels, we hypothesize that *Cartpt* may additionally be regulated by other distinct transcription factors and/or HPTMs following cocaine. This is supported by our correlational data that shows minimal overlap in the transcriptomic and epigenetic profiles of *Nr4a1* and *Cartpt* in either drug conditions in males or females in the three regions examined.

Based on these findings and the literature, we hypothesize that additional HPTMs, such as acetylation, regulate expression of *Nr4a1* and *Cartpt*. Intracranial HPC injection of the histone deacetylase (HDAC) inhibitor, Trichostatin A, increases *Nr4a1* mRNA and protein levels (48), as does HDAC3 inhibition in the dorsal HPC (49). In cell lines, *Nr4a1* expression is regulated by the recruitment of the histone acetyltransferase p300 or HDAC1 (50). It is interesting to consider that changes in STR histone acetylation, and methylation to a certain extent, are sensitive to the duration of cocaine treatment (acute versus chronic) and time since the last injection (11), which may reflect the rapid turnover of acetylation, especially compared to methylation (39). Although acetylation turnover is more rapid than methylation, we hypothesize that if we repeated our experiment at a different time point, such as 30 minutes after the final injection versus 24 hours, we would likely see different levels of H3K4me3, H3K27me3, and K4&K27 bivalency. Specifically, we expect an initial decrease in H3K27me3 and K4&K27 bivalency (<30 minutes following cocaine) followed by an increase in H3K27me3 and K4&K27 bivalency at later timepoints (>24 hours following cocaine).

Nonetheless, our findings encourage future studies to investigate combinatorial HPTMs beyond K4&K27 bivalency. For example, H3K27me3 and H3K27ac bivalency are well described in the literature and represent a tractable starting point for further exploration (51,52). We hypothesize that cocaine may also induce deenrichment of H3K27ac at *Nr4a1* and subsequently, inhibit the activation of *Nr4a1* expression in males. Finally, in the current study we test and support the hypothesis that K4&K27 bivalency is increased at *Nr4a1* in male mice following cocaine treatment, concomitant with increased H3K27me3 enrichment. To test the sufficiency of loss of H3K27me3 in increased *Nr4a1* mRNA, we can apply *in vivo* epigenetic editing to exogenously enrich *Nr4a1* H3K27me3 in the presence of cocaine. This approach may also shed light onto the role of H3K27ac, given that dCas9-FOG1 interacts with the NuRD complex to cause HDAC1/2-mediated removal of H3K27ac (53).

In closing, despite epidemiological evidence that cocaine abuse afflicts both men and women (54), the underlying neurobiology of how cocaine impacts each sex is not fully understood. This is problematic as cocaine causes distinct neuroadaptations for males and females (55). Additionally, women have a quicker escalation to drug abuse (56) and are more vulnerable to cue-induced cocaine craving compared to men (57). Previous studies also report that female rodents are more responsive to drug-conditioned stimuli compared to male rodents (58,59). The inclusion of both sexes within research investigating reward neurobiology is vital and necessary. Future studies should include both males and females to understand how cocaine causes sex-specific neuroadaptations and for uncovering sexually dimorphic mechanisms that could lead to innovations for therapeutic intervention.

## Acknowledgments & Disclosures

Financial support was provided by NIH-NIDA Avenir Director’s Pioneer Award (E.A.H, DP1 DA044250), NIDA Research Project Grant (E.A.H, 1R01DA052465-01A1), Pilot Award from the Epigenetics Institute at the University of Pennsylvania, T32 Predoctoral Training Grant in Addiction (D.K.F. NIDA T32DA028874), Penn Undergraduate Research & Mentoring program (C.H.) and The Endowed Dean’s Research Award at the University of Pennsylvania (C.H.). We thank Dr. Heath Schmidt (University of Pennsylvania) for his continuous support and meticulous review of the data and manuscript. Figure 1 was created with Biorender.com. The authors report no biomedical financial interests or potential conflicts of interest.

