## Supplemental Tables for "Cocaine regulation of *Nr4a1* chromatin bivalency and mRNA in male and female mice"

| Promoter Region | Forward Sequence | Reverse Sequence |
|---|---|---|
| Nr4a1 | ATTTACAACACCCCCTCCTCC | TTCCATTGACGCAGGGAGCG |
| Cartpt | ACACAAGAGCCGTCAATTCCA | TCGAGTTCCCAACACCGC |

**Supplementary Table 1.** Primer sequences used in qChIP.

| Gene Name | Forward Sequence | Reverse Sequence |
|---|---|---|
| *Nr4a1*_Exon4/5 | ATGCCTCCCCTACCAATCTTC | CACCAGTTCCTGGAACTTGGA |
| *Nr4a1*_Exon6/7 | AGCTTGGGTGTTGATGTTCC | AATGCGATTCTGCAGCTCTT |
| *Cartpt* | ACGAGAAGGAGCTGATCGAA | TCTCTGAGGGGAACGCAAAC |
| *A2A* | ACTCTCCCCTCCACACCC | CATAGTTTCTGTCTTCCAGCCC |
| *Drd1* | TTCTTCCTGGTATGGCTTGG | GCTTAGCCCTCACGTTCTTG |
| *Drd2* | TGGACTCAACAACACAGACCAGAATG | GATATAGACCAGCAGGTTGACGATGA |
| *Gapdh* | AGGTCGGTGTGAACGGATTTG | TGTAGACCATGTAGTTGAGGTCA |

**Supplementary Table 2.** Primer sequences used in RT-qPCR.

| Sex | Region | Drug Condition | Relationship | +/- | Related Figure(s) |
|---|---|---|---|---|---|
| M | STR | Saline | *Nr4a1* mRNA (Exon 6/7) – *Nr4a1* H3K27me3/H3K4me3 | - | 2A, 2E |
| M | STR | Saline | *Nr4a1* H3K4me3 – *Nr4a1* H3K27me3 | + | 2B, 2C |
| M | STR | Saline & Cocaine | *Nr4a1* H3K4me3 – *Cartpt* H3K4me3 | + | 2B, 3B |
| M | STR | Saline & Cocaine | *Nr4a1* (Exon 4/5) – *Nr4a1* (Exon 6/7) | + | 2A |
| M | STR | Cocaine | *Cartpt* mRNA – *Cartpt* H3K27me3 | + | 3A, 3C |
| M | STR | Cocaine | *Cartpt* mRNA – *Nr4a1* H3K4me3/H3K27me3 | + | 3A, 2D |
| M | STR | Cocaine | *Cartpt* H3K27me3 – *Nr4a1* H3K4me3/H3K27me3 | + | 3C, 2D |
| F | STR | Saline | *Nr4a1* mRNA (Exon 4/5) – *Cartpt* H3K27me3 | + | 4A, 5C |
| F | STR | Saline | *Nr4a1* mRNA (Exon 6/7) – *Cartpt* H3K27me3 | + | 4A, 5C |
| F | STR | Cocaine | *Nr4a1* H3K4me3 – *Cartpt* H3K4me3 | + | 4B, 5B |
| F | STR | Cocaine | *Nr4a1* H3K27me3 – *Cartpt* H3K4me3 | + | 4C, 5B |
| F | STR | Saline & Cocaine | *Nr4a1* (Exon 4/5) – *Nr4a1* (Exon 6/7) | + | 4A |

**Supplementary Table 3.** Significant negative (-) or positive (+) relationships in the Striatum (STR) defined by Pearson's correlation matrices from RT-qPCR and qChIP data for male (M) and female (F) mice.

| Sex | Region | Drug Condition | Relationship | Corr. | Related Figure(s) |
|---|---|---|---|---|---|
| M | HPC | Saline | *Nr4a1* mRNA (Exon 4/5) – *Nr4a1* H3K27me3 | - | 2F, 2H |
| M | HPC | Saline | *Nr4a1* mRNA (Exon 4/5) – *Cartpt* H3K4me3 | - | 2F, 3E |
| M | HPC | Cocaine | *Nr4a1* (Exon 4/5) – *Nr4a1* (Exon 6/7) | + | 2F |
| M | HPC | Cocaine | *Nr4a1* mRNA (Exon 4/5) – *Cartpt* H3K4me3 | + | 2F, 3E |
| M | HPC | Cocaine | *Cartpt* mRNA – *Nr4a1* H3K27me3/H3K4me3 | + | 3D, 2J |
| F | HPC | Saline | *Nr4a1* H3K4me3 – *Cartpt* H3K4me3 | + | 4G, 5E |
| F | HPC | Saline & Cocaine | *Nr4a1* H3K27me3 – *Cartpt* H3K27me3 | + | 4H, 5F |
| F | HPC | Cocaine | *Nr4a1* (Exon 4/5) – *Nr4a1* (Exon 6/7) | + | 4F |
| F | HPC | Cocaine | *Cartpt* mRNA – *Cartpt* H3K4me3 | - | 5D, 5E |
| F | HPC | Cocaine | *Nr4a1* (Exon 4/5) – *Cartpt* H3K4me3 | + | 4F, 5E |
| F | HPC | Cocaine | *Nr4a1* H3K27me3 – *Cartpt* H3K4me3 | + | 4H, 5E |

**Supplementary Table 4.** Significant negative (-) or positive (+) relationships in the Hippocampus (HPC) defined by Pearson's Correlation Matrices from RT-qPCR and qChIP data for male (M) and female (F) mice.

| Sex | Region | Drug Condition | Relationship | Corr. | Related Figure(s) |
|---|---|---|---|---|---|
| M | PFC | Saline | *Nr4a1* mRNA (Exon 4/5) – *Nr4a1* H3K4me3/H3K27me3 | + | 2K, 2N |
| M | PFC | Saline & Cocaine | *Nr4a1* H3K4me3 – *Cartpt* H3K4me3 | + | 2L, 3H |
| M | PFC | Cocaine | *Nr4a1* H3K4me3/H3K27me3 – *Nr4a1* H3K27me3/H3K4me3 | + | 2N, 2O |
| M | PFC | Cocaine | *Nr4a1* (Exon 4/5) – *Nr4a1* (Exon 6/7) | + | 2K |
| M | PFC | Cocaine | *Nr4a1* H3K4me3/H3K27me3 – *Cartpt* H3K4me3 | + | 2N, 3H |
| M | PFC | Cocaine | *Nr4a1* H3K4me3 – *Cartpt* H3K4me3 | + | 2L, 2H |
| F | PFC | Saline | *Nr4a1* mRNA (Exon 6/7) – *Nr4a1* H3K4me3/H3K27me3 | + | 4K, 4N |
| F | PFC | Saline & Cocaine | *Nr4a1* (Exon 4/5) – *Nr4a1* (Exon 6/7) | + | 4K |
| F | PFC | Cocaine | *Nr4a1* (Exon 4/5) – *Nr4a1* H3K27me3/H3K4me3 | - | 4K, 4O |
| F | PFC | Cocaine | *Nr4a1* (Exon 6/7) – *Nr4a1* H3K27me3/H3K4me3 | - | 4K, 4O |
| F | PFC | Cocaine | *Nr4a1* (Exon 4/5) – *Cartpt* mRNA | - | 4K, 5G |
| F | PFC | Cocaine | *Cartpt* mRNA – *Nr4a1* H3K4me3/H3K27me3 | + | 5G, 4N |

**Supplementary Table 5.** Significant negative (-) or positive (+) relationships in the Prefrontal Cortex (PFC) defined by Pearson's correlation matrices from RT-qPCR and qChIP data for male (M) and female (F) mice.
